# Mitochondrial translation occurs preferentially in the peri-nuclear mitochondrial network of cultured human cells

**DOI:** 10.1101/2021.09.15.460460

**Authors:** Christin Albus, Rolando Berlinguer-Palmini, Caroline Caroline, Fiona McFarlane, Elisabeta-Ana Savu, Robert N. Lightowlers, Zofia M. Chrzanowska-Lightowlers, Matthew Zorkau

**Affiliations:** Wellcome Centre for Mitochondrial Research, Newcastle University Biosciences Institute, Faculty of Medical Sciences, Newcastle upon Tyne, NE2 4HH, UK; Newcastle University, Bioimaging Unit, Faculty of Medical Sciences, Newcastle upon Tyne, NE2 4HH, UK; Newcastle University, Newcastle upon Tyne, NE2 4HH, UK

**Author notes:** Co-correspondence; (CA and MZ)).

**Keywords:** Mitochondria, mammalian, protein synthesis, heterogeneity, co-localisation, perinuclear, peripheral

## Abstract

Human mitochondria are highly dynamic organelles, fusing and budding to maintain reticular networks throughout many cell types. Although extending to the extremities of the cell, the majority of the network is concentrated around the nucleus in most of the commonly cultured cell lines. This organelle harbours its own genome, mtDNA, with a different gene content to the nucleus, but the expression of which is critical for maintaining oxidative phosphorylation. Recent advances in click chemistry have allowed us to visualise sites of mitochondrial protein synthesis in intact cultured cells. We show that the majority of translation occurs in the peri-nuclear region of the network. Further analysis reveals that whilst there is a slight peri-nuclear enrichment in the levels of mitoribosomal protein and mitochondrial rRNA, it is not sufficient to explain this substantial heterogeneity in distribution of translation. Finally, we also show that in contrast, a mitochondrial mRNA does not show such a distinct gradient in distribution. These data suggest that the relative lack of translation in the peripheral mitochondrial network is not due to an absence of mitoribosomes or an insufficient supply of the mt-mRNA transcripts.

## 1. Introduction

Mitochondria are organelles found in virtually all eukaryotes and all nucleated cells in the human body. They are integral to numerous functions such as the intrinsic pathway of apoptosis, generating Fe-S clusters as cofactors for important proteins and housing the enzymes responsible for pathways of oxidative metabolism. Central to the latter function is the formation of five key multi-subunit enzyme complexes that together couple cellular respiration to the production of ATP. This process is termed oxidative phosphorylation (OXPHOS) and it is the terminal step in oxidative metabolism. Four of these five complexes share a unique feature in that they contain components that are encoded by the mitochondrial genome (mtDNA), which are translated within the mitochondrial matrix. Consequently, mitochondrial gene expression is essential for cell viability.

Although original studies were interpreted to suggest that mammalian mitochondria formed many discrete organelles in the cytoplasm of the cell, more recent work has shown very clearly that mitochondria in nearly all cell types form dynamic networks, fusing and budding in a process that is choreographed by a handful of well described proteins [1]. Within the network, the inner mitochondrial membrane has large numbers of infoldings or cristae that are also remarkably dynamic, extending into the matrix and harbouring the majority of the OXPHOS supercomplexes [2–4]. The inner membrane can therefore be considered as having two discrete domains, the inner boundary membrane that parallels the outer membrane and the infoldings that constitute the cristal membranes. The two are retained as distinct by cristae junctions that largely comprise the MICOS and associated proteins [5]. This internal architecture has been suggested to isolate subcompartments, such that cristae within the same section of a mitochondrial network can support disparate membrane potentials, reflecting a heterogeneous bioenergetic status within a single network [6].

Our understanding of mitochondrial gene expression has grown enormously in recent years [7, 8]. Mitochondrial DNA is condensed to form many discrete nucleoids in the network [9–11]. This relatively short (~16.5 kb) molecule is transcribed virtually in its entirety from both strands to form polycistronic RNA units that are processed and matured in RNA granules [12, 13] that are found close to the nucleoids [14, 15]. The resultant mature RNAs are then assembled into mitoribosomes (mt-rRNAs and a subset of mt-tRNA^Val^), aminoacylated (mt-tRNAs) or translated (mt-mRNAs).

Protein synthesis is essentially the last step of gene expression and the process of oxidative phosphorylation that culminates in the generation of ATP requires synthesis of all 13 proteins that are encoded in the human mtDNA [16]. The molecular machine responsible for intramitochondrial translation is the mitoribosome, which is made up of the small (mt-SSU) and large (mt-LSU) subunits comprising ~80 proteins [17] all of which are encoded by the nuclear genome, together with two mtDNA encoded ribosomal rRNAs (*RNR1* and *RNR2*) and the *mt-tRNA^Val^* that takes the place of the 5S rRNA in the large subunit [18]. Unlike for transcription, processing and maturation, the exact submitochondrial locale of actively translating mitoribosomes has, until recently, not been clear.

By exploiting the technique of click chemistry in a process termed mitochondrial FUNCAT (*F*l*u*orescent *N*on*c*anocial *A*minoacid *T*agging), our laboratory was able to show that newly synthesised proteins were mostly located at the cristal membranes [19]. We now report that further use of mitochondrial FUNCAT reveals a heterogeneous distribution of translation within the mitochondrial network in cultured cells, with a far greater level of synthesis occurring in the perinuclear region of the network as compared to the periphery after normalisation to mitochondrial mass. Quantification of mitoribosomal protein (immunofluorescence) or mt-RNAs (RNA-FISH) confirms this mitochondrial protein synthesis gradient cannot completely be explained by a level of perinuclear enrichment of mitoribosome and mt-mRNA.

## 2. Materials and Methods

### 2.1. Cell Culture

Cultured dermal human fibroblasts, U2OS or HeLa cells were grown in Dulbecco’s modified Eagle’s medium, (Sigma D6429) supplemented with 10% foetal calf serum (Sigma), 1x non-essential amino acids and 50 μg/ml uridine (Thermo Fisher Scientific) at 37°C in humidified 5% CO_2_. Access to samples that were excess to diagnostic requirements and were approved for research was covered by the license “Role of mitochondrial abnormalities in disease” (REC ref 2002/205) issued by Newcastle and North Tyneside Local Research Ethics Committee.

### 2.2. Fluorescent non-canonical amino acid tagging (FUNCAT)

For FUNCAT experiments, cells were first seeded onto glass coverslips and cultured for 1-2 days. *In vivo* labelling of mitochondrial translation products was performed essentially as described [19]. Cells were pulsed (30 min, unless otherwise stated) with methionine-free DMEM containing the methionine analogue HPG (Jena Bioscience) with concomitant inhibition of cytosolic translation by cycloheximide (50 μg/ml). Cells were fixed before performing the copper-catalysed azide–alkyne cycloaddition of Alexa Fluor 594-picolyl-azide (CLK-1296-1) in a click chemistry reaction.

### 2.3. Immunofluorescent cytochemistry

Specific mitochondrial targets of interest were co-labelled following 30 min of HPG and incorporation using primary antibodies diluted in 5% BSA for 1 hour at room temperature, followed by species-specific Alexa Fluor 532 (Thermo Fisher A11009 or A11002) or Alexa Fluor 594 (abcam ab150116 or Thermo Fisher A11012) diluted 1:200 in 5% BSA for 40 min. Nuclei were stained with Hoechst for 5 min and cells were mounted with ProLong Glass Antifade mountant. Primary antibodies used were TOM20 (Abcam, ab78547; Santa Cruz Biotechnology, SC-17764), mL45 (Protein Tech, PA5-54778), mS27 (Protein Tech, 17280-1-AP) and ATP5I (Protein Tech, 16483-1-AP).

### 2.4. RNA FISH / Immunofluorescent cytochemistry

RNA FISH/IF was carried out as in [20] with minor modifications: Permeabilisation was carried out for 10 min, followed by incubation with respective antibodies (as above) in PBS. Stellaris Custom RNA FISH probe sets for rRNA and mRNA were designed with high stringency (masking level 5 and 4) respectively using the Stellaris Designer tool (https://www.biosearchtech.com/stellaris-designer). Probe sets were purchased with either Quasar 570 or CAL Fluor 610 fluorophores from LGC Biosearch Technologies.

### 2.5. Confocal imaging

Confocal imaging was performed on a Leica TCS SP8 (Leica Microsystems) microscope equipped with white light lasers and HC PL APO 40x/1.30 Oil CS2 objective for confocal imaging were used. The fluorophore Alexa Fluor 532 was excited at 527 nm and Alexa Fluor 594 was excited at 590 nm. Images were deconvolved using Huygens software (Scientific Volume Imaging, www.svi.nl).

### 2.6. Image analysis

In each case, approximately 100 cells were analysed from a collective of three biologically repeated experiments. For characterising colocalisation between mitochondrial markers, deconvolution and pixel based colocalisation was applied (Scientific Volume Imaging). The colocalisation analysis utilised the non-parametric Spearman’s rank correlation coefficient to determine correlation between fluorophore intensities, while Manders’ colocalisation coefficients (M1) were used to calculate proportions of co-occurrence between 0 and 1, which we have described in the text as a percentage with 1 equal to 100%. The means are given with the 95% confidence intervals. To determine any significant differences between populations, the non-parametric paired Mann-Whitney U (Wilcoxon rank sum) Test was performed with a continuity correction where necessary as implemented in the wilcox.test function in R.

## 3. Results

### 3.1. Mitochondrial translation occurs preferentially in the perinuclear network

Initial data from mitochondrial FUNCAT visualising mitochondrial protein synthesis in fibroblasts, U2OS and HeLa cells had suggested an enrichment of translation in the perinuclear region of the cell compared to the periphery when normalized to a marker of mitochondrial mass [19]. To extend this analysis we subjected human cultured U2OS cells to mitochondrial FUNCAT as previously described [19, 20]. All cell images were manually masked such that signals were assigned either to the perinuclear or peripheral regions of each cell. By quantifying the perinuclear and peripheral signals from a minimum of 50 cells and normalizing the signal to the mitochondrial marker TOM20, an outer membrane component of the protein import complex, we were able to show a distinct depletion of nascent protein synthesis towards the periphery of the cell (Figure 1 A).

**Figure 1.**
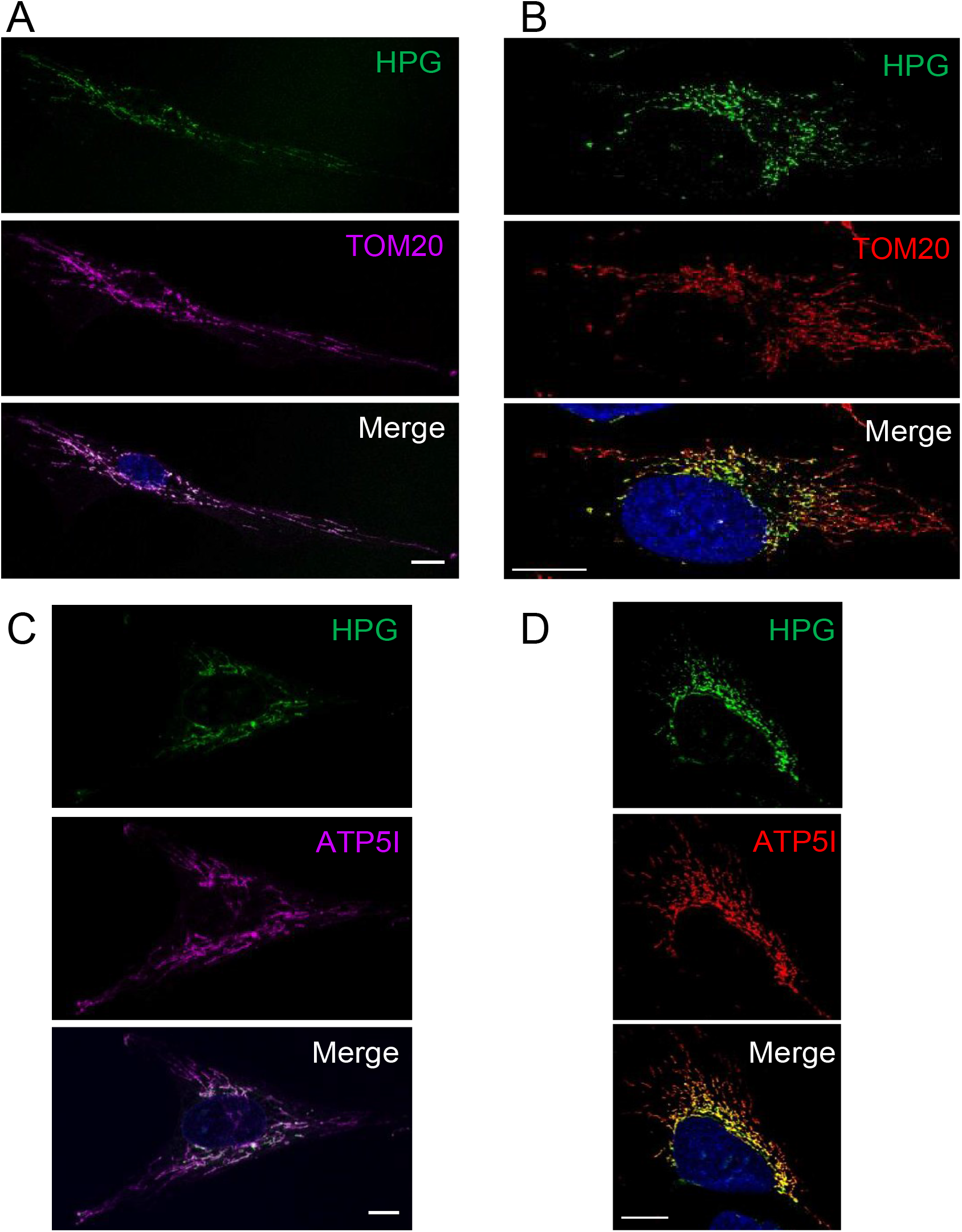
Preferential perinuclear mt-translation occurs in multiple cell types. Cells were pulsed with HPG for 30 mins in the presence of cycloheximide. Dermal fibroblast (A, C) and U2OS (B, D) cells were then fixed, click reactions performed and stained with antibodies against TOM20 (A, B) or ATP5I (C, D). Representative confocal images are shown. Scale bar = 10 μm.

We wished to determine whether this marked intracellular gradient was a simple reflection of a similar gradient of the mature OXPHOS complexes. We chose to compare the pattern of synthesis with a marker, ATP5I, of the multi-subunit FoF1 ATP synthase that contains two mtDNA encoded components and is found mainly in the mitochondrial cristae, the infoldings of the mitochondrial inner membrane [21–23]. As shown in Figure 1 B and quantified in Figure 4, a similar gradient of HPG signal diminishing towards the cell periphery is noted when compared to the immunofluorescent signal associated with ATP5I. These data confirm a strong concentration of mt-protein synthesis occurring preferentially within the perinuclear regions.

### 3.2. Preferential perinuclear translation is not reflective of a similar distribution of mitoribosomes

The most likely explanation for this gradient in new protein synthesis across the mitochondrial network (Figure 2A) could simply be defined by a similar gradient in mitoribosome localization. To address this possibility, confocal images were collected to facilitate comparison of the distribution of immunofluorescent signal of protein markers of the large (mL45, Figure 2B) and small (mS27, Figure 2C) mitoribosomal subunits. Similar normalization to TOM20, as a marker of mitochondrial mass, reveals only a modest reduction in peripheral signal compared to the perinuclear signal for each of the mitoribosomal proteins (MRPs) (5% reduction for mL45 and 10% for mS27, compared to 55% reduction for HPG representing protein synthesis). To determine whether these values are artificially high as they represent a mixture of free (unassembled) MRPs or fully assembled mitoribosomes, we subjected cells to RNA FISH with probes specifically to the ribosomal RNA components of the small (*RNR1*, Figure 2D) and large (*RNR2*, Figure 2E) mitoribosomal subunits.

**Figure 2.**
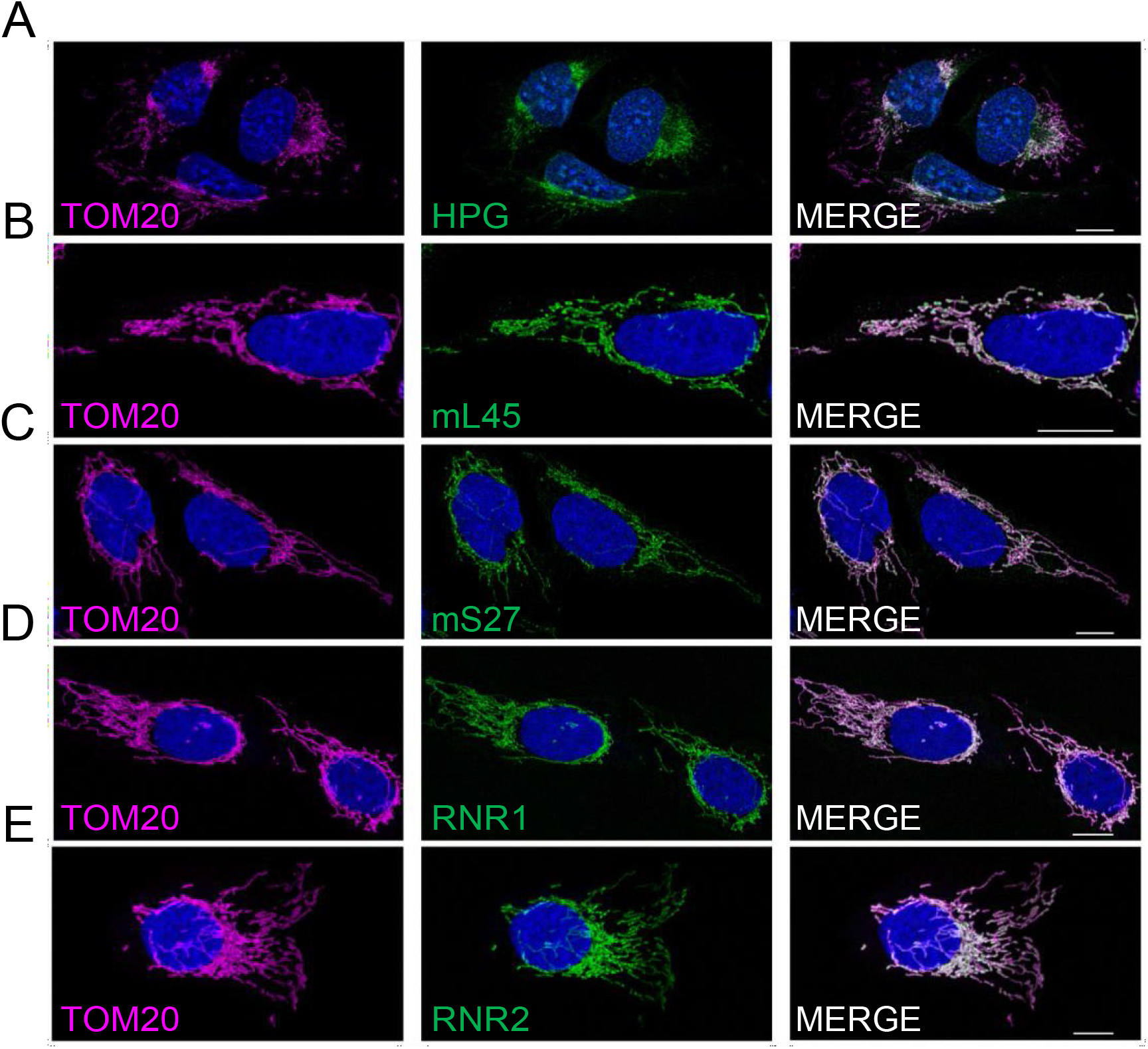
Mitoribosomal components show a different distribution to the HPG marker of mitochondrial translation. U2OS cells were immunostained with antibodies against TOM20 as a marker of the mitochondrial network either following HPG incubation and click chemistry (A) or co-stained with antibodies against mitoribosomal proteins of the large (mL45, panel B) or small (mS27, panel C) subunits. To determine distribution of the RNA components of the small (*RNR1*, panel D) and large (*RNR2*, panel E) mitoribosomal subunits, immunostaining of TOM20 was combined with RNA FISH. Representative confocal images are shown. All scale bars = 10.01 μm.

Quantification of images showed a similar reduction in the peripheral signal compared to the perinuclear signal for both the RNA and protein components of the small subunit (*RNR1* 18%, mS27 10%). However, a relatively greater reduction in signal was noted for the rRNA component of the mt-LSU, *RNR2* (26%) compared to mL45 protein (5%). As it is highly likely that ribosomal RNA will not be stable in the absence of bound mitoribosomal proteins, the more marked reduction of the *RNR2* signal in the periphery as compared to a mt-LSU protein marker could reflect the presence of a subset of free mL45 towards the periphery of the cell. Crucially, however, the decrease in peripheral signal noted for *RNR2* is still substantially less marked than for the HPG signal (26% decrease for *RNR2* cf. 55% decrease for HPG). Therefore, the extensive depletion in new protein synthesis that occurs in the peripheral mitochondrial network cannot be explained merely by a similar loss of mitoribosomes.

### 3.3. Differential distribution of mt-protein synthesis is not reflective of the mt-mRNA distribution

As the diminished level of mitochondrial translation occurring in the peripheral mitochondrial network is not due to a paralleled loss of mitoribosomes, we next determined whether it could be explained by a lack of mt-mRNA available to load onto the mitoribosomes.

A similar protocol that had been used to visualize mitoribosomal rRNA by RNA FISH [20] was employed to detect the mt-mRNA that encodes COXII of complex IV. As shown in Figure 3A, once again there was a noticeable decrease in signal towards the periphery of the cell (35%) even after normalization to the mitochondrial network marker TOM20. This depletion, however, was not comparable to the more extensive loss in distribution of the HPG signal (35% cf. 55% Figure 3A vs Figure 2A).

**Figure 3.**
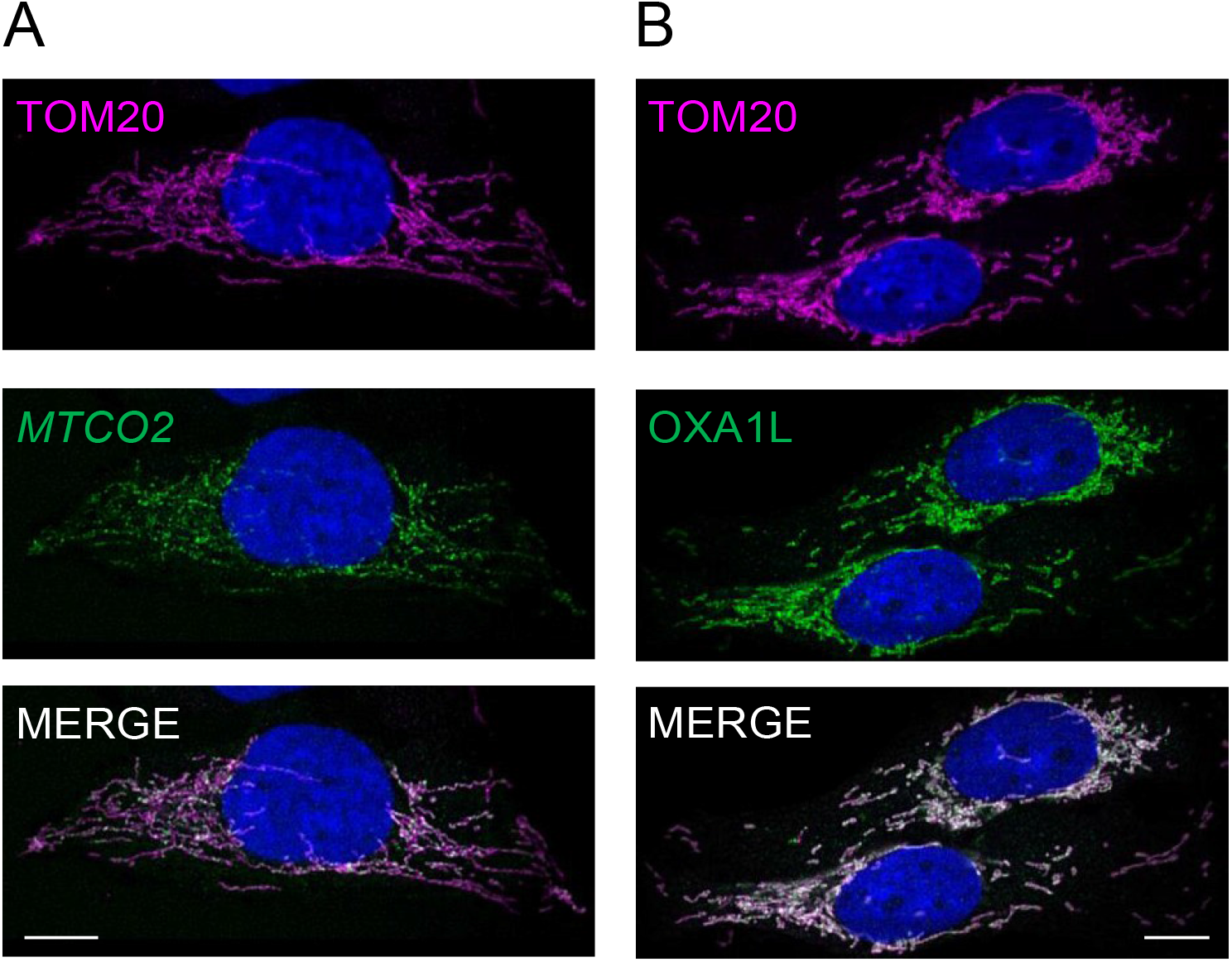
Neither transcript nor insertase levels appear to be limiting factors for mt-translation in peripheral regions of the reticulum. U2OS cells were immunostained with TOM20 antibodies to delineate the mitochondrial network either in combination with RNA FISH to define distribution of the mtDNA encoded *MTCO2* transcript (A) or co-stained with antibodies against the insertase OXA1L (B). Representative confocal images are shown. All scale bars = 10.06 μm.

### 3.4. Peripheral depletion of mitochondrial translation cannot be explained by the distribution of OXAL1

Although based solely on the use of the *MTCO2* RNA probe as an indicator of mt-mRNAs, it can be tentatively assumed that the decrease in translation is not correlated with a matching decrease in either mitoribosomes or in the availability of mt-mRNAs. What could be responsible for such a marked loss of translation in the peripheral mitochondrial network ? The inner mitochondrial membrane insertase, OXAL1 has for several years been implicated as a key player in facilitating insertion of the highly hydrophobic proteins encoded by mtDNA [24]. A more recent cryoEM study revealed how interactions between OXA1L and mL45 in the assembled mitoribosomes are able to couple protein synthesis to the insertion of the newly made protein into the inner membrane [25, 26]. We therefore reasoned that the presence of OXA1L may dictate the distribution of newly translated protein. To determine whether or not the localization of OXA1L was homogenous throughout the mitochondrial reticulum, we performed immunofluorescence with antibodies against OXA1L and TOM20 (Figure 3B).

Masking and quantification of these confocal images revealed that whilst the level of signal for OXA1L is reduced towards the periphery of the cell (28%), again it does not accurately track the loss of HPG signal (55%; Figure 4), establishing that this marked depletion in protein synthesis could not be completely due to a lack of OXA1L.

**Figure 4.**
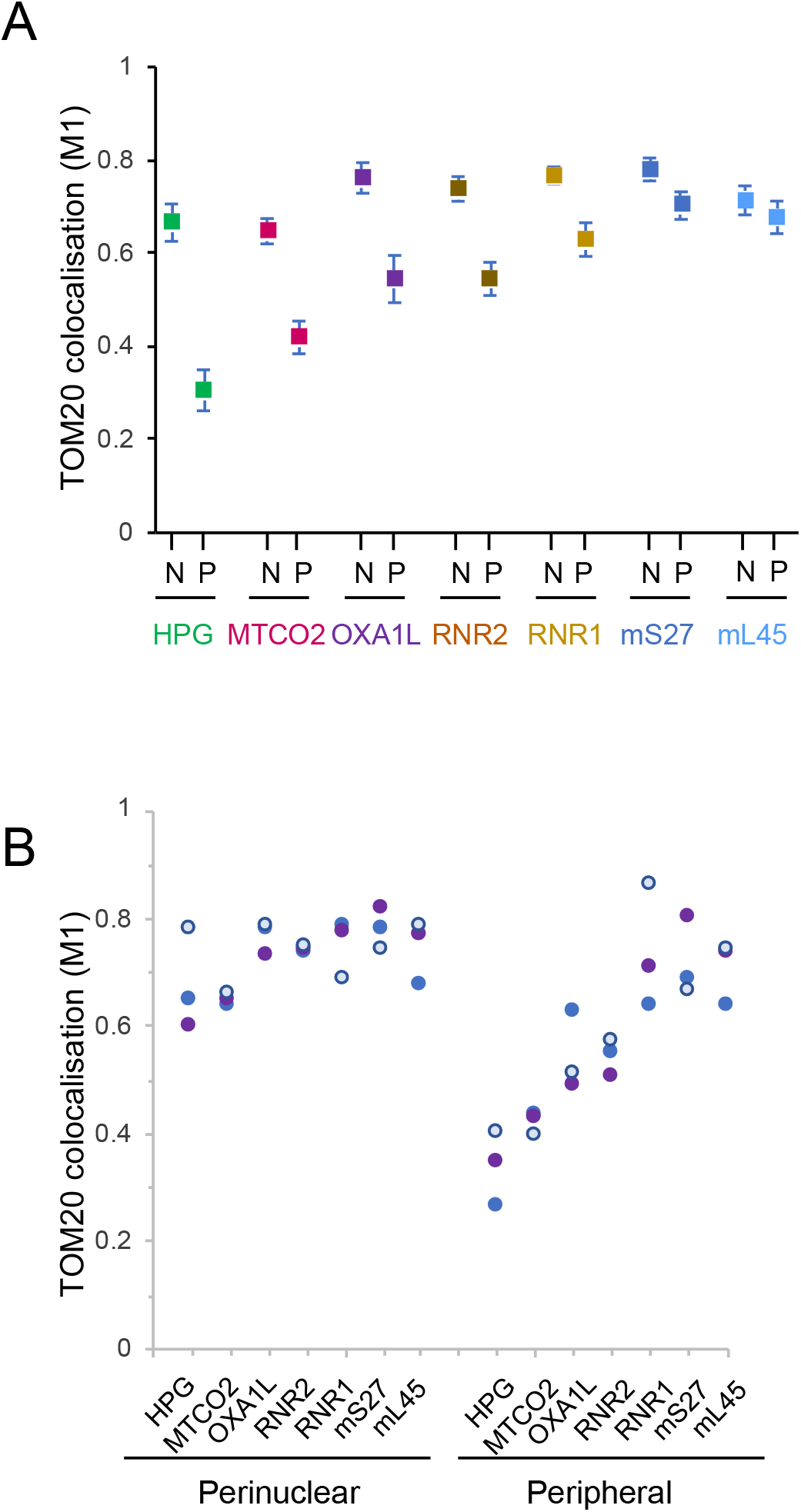
There is a differential distribution within the network of components involved in mitochondrial translation. In each case a minimum of 50 cells were analysed across 3 biological repeat experiments. Following deconvolution the correlation between fluorophore intensities of HPG (n = 54), *MTCO2* (n = 50), *RNR1* (n = 72), *RNR2* (n = 77), mS27 (n = 59), mL45 (n = 54), Oxa1L (n = 51) normalized to TOM20 in both the perinuclear and peripheral regions was calculated using the Manders’ colocalisation coefficient (M1) to determine relative co-occurrence. The mean and 95% confidence intervals are shown (A). The means (blue, white and purple circles) for each parameter are plotted to show the distribution of signal in the perinuclear versus peripheral regions for three biological repeats (B).

## 4. Discussion

The focus of this mini-series is mitochondrial heterogeneity. Our data presented here and that of another recent publication [19] show clearly that for at least certain human cell lines there is evidence of marked heterogeneity in the location of translation within the mitochondrial network. Normalising mitochondrial signal using markers to either the outer mitochondrial membrane (TOM20) or the cristae membranes (FoF1 ATP synthase) confirm new protein synthesis is more markedly concentrated within the perinuclear mitochondrial network rather than towards the cell periphery. Interestingly, although we have normalised quantification of markers to the outer membrane marker TOM20, it is also known to show a gradient itself within the network, although clearly this is not as dramatic as the translation gradient [27]. A similar perinuclear enrichment had been originally claimed for mtDNA replication [28] but further studies involving careful normalisation to mitochondrial mass showed this was not the case [28, 29]. To ensure our observations are robust, we have mapped the translation machinery through both the rRNA and protein components of the mitoribosome. We have looked at the localisation of a transcript (*MTCO2*) that is to be translated and the protein (OXA1L) that facilitates insertion of the newly synthesised proteins into the inner mitochondrial membrane. Extensive colocalisation studies now show that this geographical heterogeneity of mt-protein synthesis is not the consequence of a similar distribution of mitoribosomal components nor of the localisation of mitochondrial mRNA, as the pattern for all these elements remains distinct from that of the HPG signal, which reflects the site of synthesis of mtDNA encoded proteins (Figure 4).

It is unclear what mechanism is responsible for this heterogeneity, or what might stimulate such variation. Perhaps the most likely explanation is that a gradient in the concentration of a key component(s) of mitochondrial translation is present across the cell or within the organelle. Due to the relatively rapid equilibration of soluble proteins it is more difficult, although not impossible, to envision how such gradients of soluble factors involved in translation could be established. Thus, the integral membrane location of OXA1L together with its crucial membrane insertase role in mitochondrial protein synthesis made this a strong candidate. Although its partial peripheral depletion may explain part of the intracellular heterogeneity in mitochondrial translation it clearly cannot be the main underlying cause. What is the significance of such a gradient ? A functional variation in the perinuclear and peripheral mitochondrial subpopulations was initially reported for the mitochondrial membrane potential, with the peripheral mitochondria being on average more hyperpolarised than the perinuclear fraction [30]. Consistent with this variation, peripheral mitochondria in several cell lines were shown to more rapidly clear agonist evoked Ca^2+^ flux from the cytosol [30]. Indeed, there are many examples of functional heterogeneity of mitochondria with respect to mitochondrial redox state, membrane potential, respiratory activity, uncoupling protein targeting and mitochondrial ROS production in cells both *in vitro* and *in vivo* (for review, see [31]). Nuclear gene expression has also been suggested to be modified by accumulation of perinuclear mitochondria [32]. One additional possibility has recently been suggested by Lu *et al*. working with rabbit cardiac myocytes; the authors detailed a variety of functions that were markedly different in populations of subsarcolemmal, intrafibrillar or perinuclear mitochondria such as sensitivity to oxidative stress or maximal Ca^2+^ uptake [33]. Intriguingly, they also report a more rapid appearance of a mitochondrially-targeted GFP marker in the perinuclear compared to the other subcellular mitochondrial populations following viral transduction. This led to the suggestion that mitochondrial biogenesis may occur more readily in perinuclear mitochondria. A logical extension would then be that new mitochondrial protein would then move out towards the periphery, where mitochondria would potentially be more stable. This model would be consistent with the observation that cells devoid of KIF5B, a key component in the intracellular movement of mitochondria, lose their peripheral mitochondria, which are suggested to be made via a process of dynamic tubulation of the perinuclear network [34]. Whilst it is a substantial extrapolation from data derived from cardiac myocytes to common human cell lines, the hypothesis that graded mitochondrial translation supports a model of graded mitochondrial biogenesis is intriguing and we expect to be able to test this soon.

## Author Contributions

Conceptualization, Z.C-L., C.A., M.Z., and R.L.; Methodology, Z.C-L., C.A., M.Z., R.B-P., and R.L.; Formal analysis, Z.C-L., C.A., M.Z., and R.L.; Investigation, C.A., M.Z., C.H., F.M., E.S. and R.B-P.; Writing—original draft preparation, Z.C-L., and R.L.; Writing—review and editing, Z.C-L., C.A., M.Z., and R.L.; Visualization, Z.C-L., C.A., M.Z., and R.L.; Supervision, Z.C-L., C.A., M.Z., and R.L.; Project administration, Z.C-L., and R.L.; Funding acquisition, Z.C-L., and R.L. All authors have read and agreed to the published version of the manuscript.

## Ethics Statements

Access to samples that were excess to diagnostic requirements and were approved for research was covered by the license “Role of mitochondrial abnormalities in disease” (REC ref 2002/205) issued by Newcastle and North Tyneside Local Research Ethics Committee.

